# Acute Aerobic Exercise Enhances Associative Learning in Active but not Sedentary Individuals

**DOI:** 10.1101/2024.10.28.620014

**Authors:** Kayleigh D. Gultig, Cornelis P. Boele, Lotte E.M. Roggeveen, Emily Soong, Sebastiaan K.E. Koekkoek, Chris I. De Zeeuw, Henk-Jan Boele

## Abstract

**Introduction:** Physical exercise has repeatedly been reported to have advantageous effects on brain functions, including learning and memory formation. However, objective tools to measure such effects are often lacking. Eyeblink conditioning is a well-characterised method for studying the neural basis of associative learning. As such, this paradigm has potential as a tool to assess to what extent exercise affects one of the most basic forms of learning. Until recently, however, using this paradigm for testing human subjects in their daily life was technically challenging. As a consequence, no studies have investigated how exercise affects eyeblink conditioning in humans. Here we hypothesize that acute aerobic exercise is associated with improved performance in eyeblink conditioning. Furthermore, we explored whether the effects of exercise differed for people with an active versus a sedentary lifestyle.

**Methods:** We conducted a case-control study using a smartphone-based platform for conducting neurometric eyeblink conditioning in healthy adults aged between 18 – 40 years (n = 36). Groups were matched on age, sex, and education level. Our primary outcome measures included the amplitude and timing of conditioned eyelid responses over the course of eyeblink training. As a secondary measure, we studied the amplitude of the unconditioned responses.

**Results:** Acute exercise significantly enhanced the acquisition of conditioned eyelid responses; however, this effect was only true for individuals with an active lifestyle. No statistically significant effects were established for timing of the conditioned responses and amplitude of the unconditioned responses.

**Discussion:** This study highlights a facilitative role of acute aerobic exercise in associative learning and emphasises the importance of accounting for lifestyle when investigating the acute effects of exercise on brain functioning.

## 1 INTRODUCTION

Physical exercise is often proposed to have beneficial effects on brain function, including learning and memory formation(1,2). The reported short- and long-term effects of ‘sports on the brain’ are, however, variable(3–5), with some suggesting the benefits to be exaggerated(6). A more objective way to investigate if and how physical activity impacts learning is through Pavlovian eyeblink conditioning, a well-established paradigm to study associative motor learning. In eyeblink conditioning, an unconditional stimulus (US) that reliably evokes a reflexive eyeblink, is repeatedly paired with a conditioned stimulus (CS). Eventually the CS itself will evoke an anticipatory eyeblink, which is called a conditioned response (CR)(7). The neural circuits and plasticity mechanisms underlying eyeblink conditioning have been studied extensively in both experimental animals and human participants. In mice, acute physical activity during eyeblink conditioning, such as voluntary or externally imposed treadmill running, enhances learning and expression of conditioned eyelid responses(8,9). However, chronic physical activity in the form of a physically enriched environment does not seem to improve learning or CR expression in rodents (10), though it has small but significant effects on the adaptive timing of CRs. To our knowledge, the effects of exercise on eyeblink conditioning in humans have not yet been investigated.

In this exploratory study, we examine the effects of acute aerobic exercise on cerebellar associative learning in healthy adults, using a smartphone-based platform to conduct eyeblink conditioning tests. We assess the effects of acute exercise in individuals with active or sedentary lifestyles. Based on the reported acute effects of physical activity in mice, and the fact that eyeblink conditioning mechanisms are conserved across species (11), we hypothesize that aerobic exercise will similarly facilitate eyeblink conditioning in humans. Furthermore, since long-term exercise influences the brain’s response to acute exercise (3), we expect that the exercise-enhancing effects on eyeblink conditioning will be greater in regularly active individuals compared to sedentary participants.

## 2 METHODS

### 2.1 Participants

40 Neurotypical participants aged between 18 and 40 years, were recruited by social media invitations to participate in the study. This sample size in this pilot study is in-line with other eyeblink conditioning research in humans (12,13). Participants were divided into an active or sedentary lifestyle group based on their weekly hours of physical activity. The cut-off point for group classification was determined using the lower limit of the WHO guidelines for physical activity in adults aged 18-64 years(14). Participants doing less than 2.5 hours of moderate intensity or less than 75 minutes of vigorous intensity exercise were in the sedentary group and the other participants were in the active group. Moderate intensity was defined as: “Exercise that increases heart rate but you are still able to hold a conversation” and vigorous as “Exercise that raises your heart rate so that you are unable to speak”. Education level was similar across groups as all subjects either had a university degree or were university students. Furthermore, the average age and hours of sleep per night were similar across groups (**table 1**). This study was approved by the Institutional Review Board for Human Subjects of Princeton University (IRB #13943). All participants were informed of the study protocol and gave their written informed consent.

**Table 1.** Demographic and exercise-related data for participants in the active and sedentary lifestyle groups with or without an exercise intervention.

|  | Active,<br>post-exercise<br>(n = 9) | Active,<br>no-exercise<br>(n=9) | Sedentary,<br>post-exercise<br>(n=9) | Sedentary,<br>no-exercise<br>(n=9) | Statistical<br>test | p-value |
| --- | --- | --- | --- | --- | --- | --- |
|  | Mean (±SD) |  |  |  |  |  |
| Moderate<br>exercise<br>(hours/week) | 3.2 (±1.6) | 4.7 (±2.4) | 0.9 (±0.8) | 1.1 (±0.8) | F<sub>3,32</sub> = 12.5 | < 0.0001 |
| Vigorous<br>exercise<br>(hours/week) | 4.1 (±3.1) | 2.1 (±1.3) | 0.2 (±0.3) | 0.0 (±0.0) | F<sub>3,29</sub> = 9.6 | 0.0001 |
| Total<br>exercise<br>(hours/week) | 7.3 (±3.4) | 6.8 (±2.9) | 1.1 (±0.8) | 1.1 (±0.8) | F<sub>3,32</sub> = 20.3 | < 0.0001 |
| Age (years) | 24.9 (±2.3) | 25.6 (±5.9) | 22.2 (±3.6) | 24.9 (±5.0) | F<sub>3,32</sub> = 1.01 | 0.40 |
| Sleep<br>(hours/night) | 7.5 (±0.8) | 7.9 (±0.3) | 7.6 (±1.2) | 7.3 (±0.4) | F<sub>3,30</sub> = 0.8 | 0.49 |
| Average HR<br>during<br>exercise<br>(bpm) | 143.7<br>(±18.6) | - | 152.4<br>(±20.0) | - | t<sub>11</sub> = -0.8 | 0.43 |
| Average<br>duration of<br>exercise<br>intervention<br>(minutes) | 33.5 (±9.6) | - | 31.0 (±3.6) | - | Z = -0.3 | 0.82 |
| Self-rated<br>exercise<br>intensity /5 | 2.8 (±0.5) | - | 3.7 (±1.0) | - | t<sub>11,51</sub> = -2.3 | 0.04 |
| Sex |  |  |  |  |  |  |
| Male | 4 (44.4%) | 4 (44.4%) | 3 (33.3%) | 5 (55.6%) | - | - |
| Female | 5 (55.6%) | 5 (55.6%) | 6 (66.7%) | 4 (44.4%) | - | - |

Statistical comparisons done using one-way ANOVAs for exercise, age and sleep; a t-test for unequal variances for exercise intensity ratings, a t-test for average heart rate and Wilcoxon's rank sum test for average exercise intervention duration; SD = Standard Deviation, HR = Heart Rate, bpm = beats per minute

### 2.2 Experiments

Experiments were conducted via the BlinkLab smartphone application(15). During the experiment, participants watched audio-normalized nature documentaries or TV shows. A delay eyeblink conditioning paradigm, a form of cerebellar associative learning(8,16), was used in this study. The eyeblink conditioning experiment consisted of the pairing of a CS with a US (previously described by Boele and colleagues(15) (**figure 1A, B**)). The CS, a circular white dot 1cm in diameter, was presented in the center of the phone screen for 450 ms. The US was a simultaneous full-screen flash and 105dB white noise pulse presented for 50ms. In paired trials, the US was presented 400 ms after the onset of the CS and co-terminated with the CS. In US-only trials, the stimuli were presented for 50 ms, 400 ms from trial onset. Each eyeblink conditioning session consisted of 10 blocks. Within each block, there were 8 paired trials, 1 CS-only trial and 1 US-only trial semi-randomly distributed throughout the block.

**Figure 1.**
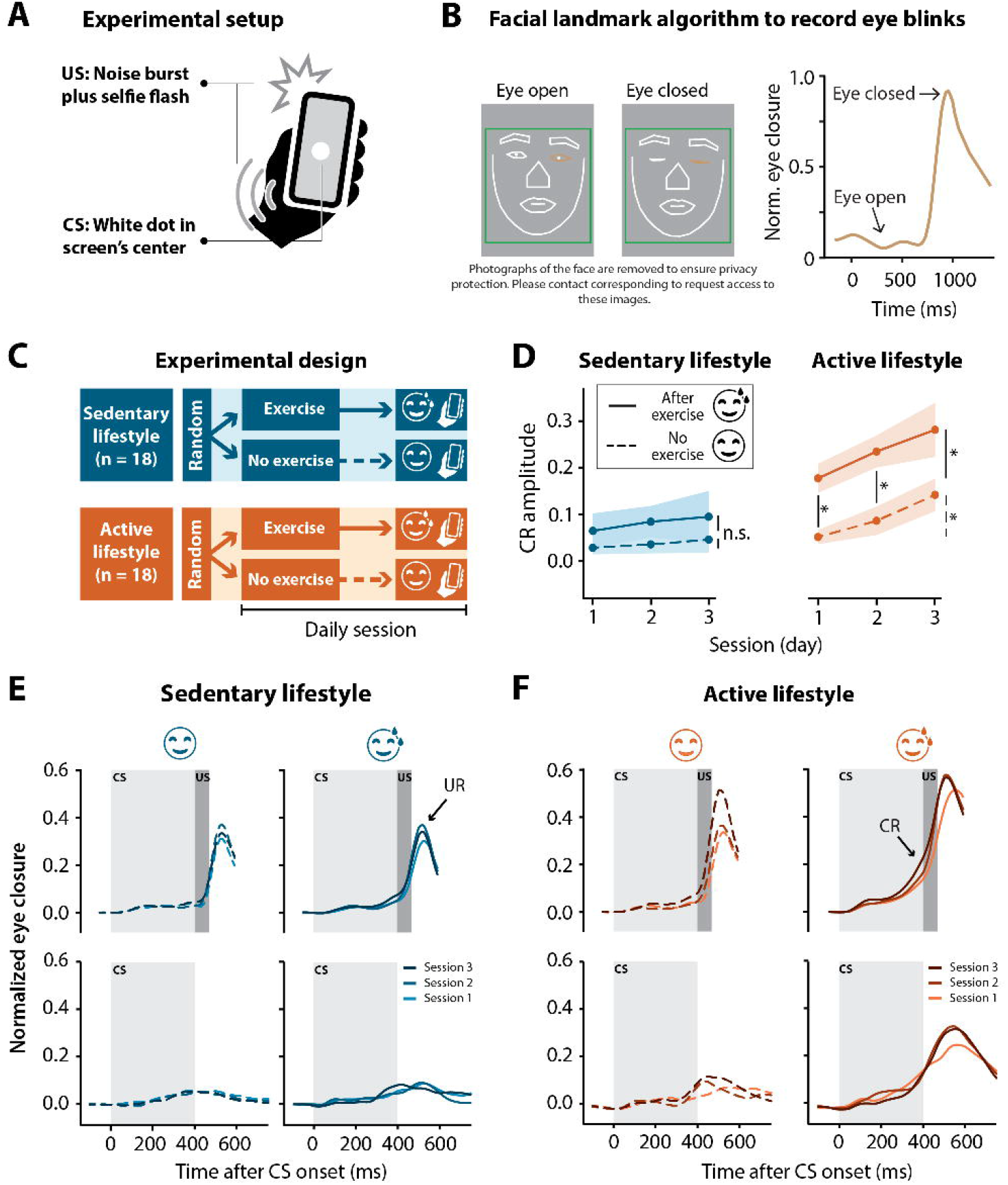
Smartphone-mediated eyeblink conditioning in participants with an active versus sedentary lifestyle. Half exercised prior to the conditioning sessions and the other half did not. **(A)** Experimental setup for smartphone-mediated eyeblink conditioning. The conditioned stimulus (CS) was a 1 cm diameter white circle presented in the center of the screen for 450 ms. The unconditioned stimulus (US) was a simultaneous 50 ms full screen flash and 105 dB 50 ms white noise pulse, presented at 400 ms and co-terminating with the CS at 450 ms. **(B)** Facial landmark detection algorithm detects eyelid movements recorded by the smartphone’s forward-facing camera in real-time. In this example, the raw signal amplitude is recorded from the left eye. A value of 0 corresponds to eye open and a value of 1 corresponds to eye closed. **(C)** Illustration of experimental design. Sedentary (< 2.5 hours of exercise in a week) and active (> 2.5 hours of exercise in a week) individuals were randomly assigned to a no-exercise or an exercise intervention. The exercise intervention completed three eyeblink conditioning sessions directly after moderate intensity running or cycling whereas the no-exercise intervention did no exercise for at least 8 hours prior to the three eyeblink conditioning sessions. No more than one session was done on a day. **(D)** Conditioned response (CR) amplitude by session for paired (CS + US) and CS-only trials combined in the sedentary and active groups with or without an exercise intervention before eyeblink conditioning sessions. Active individuals showed significant conditioning with the post-exercise intervention showing significantly higher conditioned response amplitudes at sessions 1 and 2 compared to the no exercise intervention. Colored shading represents standard error of the mean. **(E)** Sedentary and **(F)** active group averaged eyelid traces for paired (CS + US) trials (top panels) and CS-only trials (bottom panels) without (left panels) or after (right panels) exercise for three eyeblink conditioning sessions. Light grey blocks indicate the presentation of the CS for 450 ms and dark grey blocks indicate the presentation of the US for 50 ms co-terminating with the CS at 450 ms. In paired trials, note the peak in amplitude following the presentation of the US, namely the unconditioned response (UR) present in all groups regardless of the intervention. Note the shift in timing of the rise in amplitude in paired trials to precede the presentation of the US at later sessions - CR - especially obvious in the active, post-exercise intervention. The acquisition of conditioned responses over the three sessions is also illustrated by the rise in amplitude in the CS-only trials, again particularly obvious in the active, post-exercise intervention. Significance levels: * p < 0.05, n.s. = not significant.

### 2.3 Procedure

#### 2.3.1 Experimental setup

Participants were instructed to use headphones and complete the experiments in a quiet, well-lit room. All participants completed three sessions of experiments in the space of a week, with no sessions done on the same day.

#### 2.3.2 Exercise groups

Participants in the active and sedentary groups were randomly assigned to an exercise and no-exercise ‘intervention’. Participants in the exercise intervention were instructed to do all eyeblink conditioning sessions as soon as possible after at least 30 minutes of moderate intensity running or cycling and to record the activity using a smartwatch if they had access to such a device. Participants in the no-exercise intervention were instructed to refrain from exercise for at least 8 hours before the test (**figure 1C**). Before starting the eyeblink training session, participants were asked, in the app, to rate the intensity of the exercise on a five-point Likert type scale(17) (**table 1**).

### 2.4 Data processing

Data processing was done in R 4.3.1. Trials were baseline corrected using the 500 ms stimulus-free baseline and min-max normalized using spontaneous blinks as a reference. Individual eyelid traces were normalized by dividing each trace by the maximum signal amplitude of the relevant session. Thus, eyes closed corresponded to a value of 1 and eyes open to a value of 0 (**figure 1B**)(15).

Trials with extreme outliers (signal amplitude < -0.4) and trials where spontaneous blinks occurred within a time window of 150 ms before, until 35 ms after stimulus presentation were excluded from further analysis. Trials were then re-baseline corrected using the same time window that was used for removal of spontaneous blinks.

CR amplitude was determined as the maximum signal amplitude value at 430 ms, for paired and CS-only trials. This time value was chosen to allow for a latency of 30ms following the expected presentation of the US at 400 ms. There is a latency in response to the US (**supplemental figure 1**) likely due to retinal processing of the flash(18).

To compare latency to CR peak between groups, CS-only trials were analyzed. Here, CRs were defined as trials with a maximum signal amplitude above 0.10 in a time window ranging from 60 – 750 ms. Additionally, the mean percentage of well-timed CRs was calculated per group. A well-timed CR was defined as a trial with a maximum signal amplitude above 0.10 in a time window between 400 – 500 ms.

### 2.5 Statistical analysis

All statistical analyses and visualizations were done in R 4.3.1. Potential differences between groups in age, average weekly exercise and sleep hours were tested using a one-way ANOVA. A t-test for unequal variances was used to compare the self-reported exercise intensity levels between the active and sedentary groups who completed eyeblink conditioning after exercise. Average heart rate was compared between active and sedentary post-exercise groups using a t-test. Wilcoxon’s rank sum test was used to compare average exercise duration for the active and sedentary groups post-exercise intervention.

For all other analyses, multilevel linear mixed effects (LME) models were used. These models are robust to deviations from normality and are more appropriate for the nested data structure of this study (19,20). In all models, ‘subject’ was used as a random effect. For CR amplitude models, a random slope for the effect of sessions across subjects was used. Data for the CR amplitude models was normalised using ordered quantile normalisation, suggested by the bestNormalize package in R, to allow for optimal model fit. Fixed effects included: ‘session’, ‘exercise’ and ‘exercise*session’. Fixed effects for the latency to CR peak models included: ‘exercise’ and ‘CR amplitude’. The models for well-timed CRs had ‘exercise’ as a fixed effect. The restricted maximum likelihood method was used to estimate model parameters. Log likelihood ratio and AIC and BIC indices were used to assess the model fit. An alpha value of p < 0.05 (two-tailed) was used to determine significance. For multiple comparisons, Bonferroni-Holm p-value adjustments were made to account for the number of comparisons.

## 3 RESULTS

### 3.1 Participant overview

A total of 40 neurotypical participants were initially included in the study, which was conducted over the period from 28 December 2022 until 31 May 2023. Three participants were excluded during data pre-processing due to hardware-related latency issues. One participant in the active, post-exercise group was excluded due to a complete lack of eyeblink startle responses in all sessions; four sessions from different participants were excluded due to WIFI-related technical issues with the application: session 1 for one participant in the sedentary, post-exercise intervention; session 1 for one participant in the active, no-exercise intervention; session 2 for one participant in the active, no-exercise intervention and session 3 for one participant in the active, no-exercise intervention. The final cohort included 18 individuals in each lifestyle group, split into nine individuals per intervention (exercise vs. no exercise) (**table 1**). The total, moderate and vigorous hours of exercise per week differed significantly between the active and sedentary groups **(table 1**). All mean values presented below are ± standard deviation (SD) and p-values are Bonferroni-Holm corrected for multiple comparisons.

### 3.2 Physical activity

Participants in the sedentary, post-exercise intervention completed all three sessions on average 11 minutes following exercise while those in the active, post-exercise intervention completed all three sessions on average 14 minutes following exercise. Two participants in the sedentary post-exercise intervention and two in the active post-exercise intervention did not perform the instructed exercise type for one of the three sessions. The frequency of exercise types completed prior to eyeblink conditioning can be seen in **supplemental figure 2**. The average duration of exercise completed prior to eyeblink conditioning can be seen in **table 1** and did not differ significantly between the active and sedentary groups (Z = -0.3, p = 0.82). Two participants in the sedentary post-exercise intervention and one in the active post-exercise intervention were not able to record their heart rate with a smartwatch. Average heart rate data for the remaining participants can be seen in table 1 and did not differ significantly between active and sedentary participants (t_11_ = -0.8, p = 0.43). Both active and sedentary individuals in the no-exercise intervention completed all three sessions without any aerobic exercise for at least 8 hours before the test.

### 3.3 Conditioning – acquisition

While some participants did not acquire CRs (**supplemental figure 3A**), in others the acquisition of CRs already started to occur in session 1 with the amplitude and timing of these responses improving over the course of three sessions (**supplemental figure 3B**).

In order to assess the effect of aerobic exercise on eyeblink conditioning we compared the CR amplitude between no and post-exercise interventions within the sedentary and active groups separately. For the sedentary group, CR amplitudes in the no exercise intervention were low (session 1 mean=0.03 ±0.15, **table 2**) and did not really increase by session 3 (mean=0.05 ±0.18). In the post-exercise intervention, CR amplitudes were slightly higher (session 1 mean=0.07 ± 0.20) and showed a slight increase by session 3 (mean=0.10 ±0.23). Within the sedentary group, no main effect of session (F_2,3476_=1.41, p=0.24) or exercise (F_1,16_=1.49, p=0.24) was found and there was no significant interaction between session and exercise (F_2,3476_=0.78, p=0.46) **figure 1D, E, supplemental table 1**).

**Table 2.**
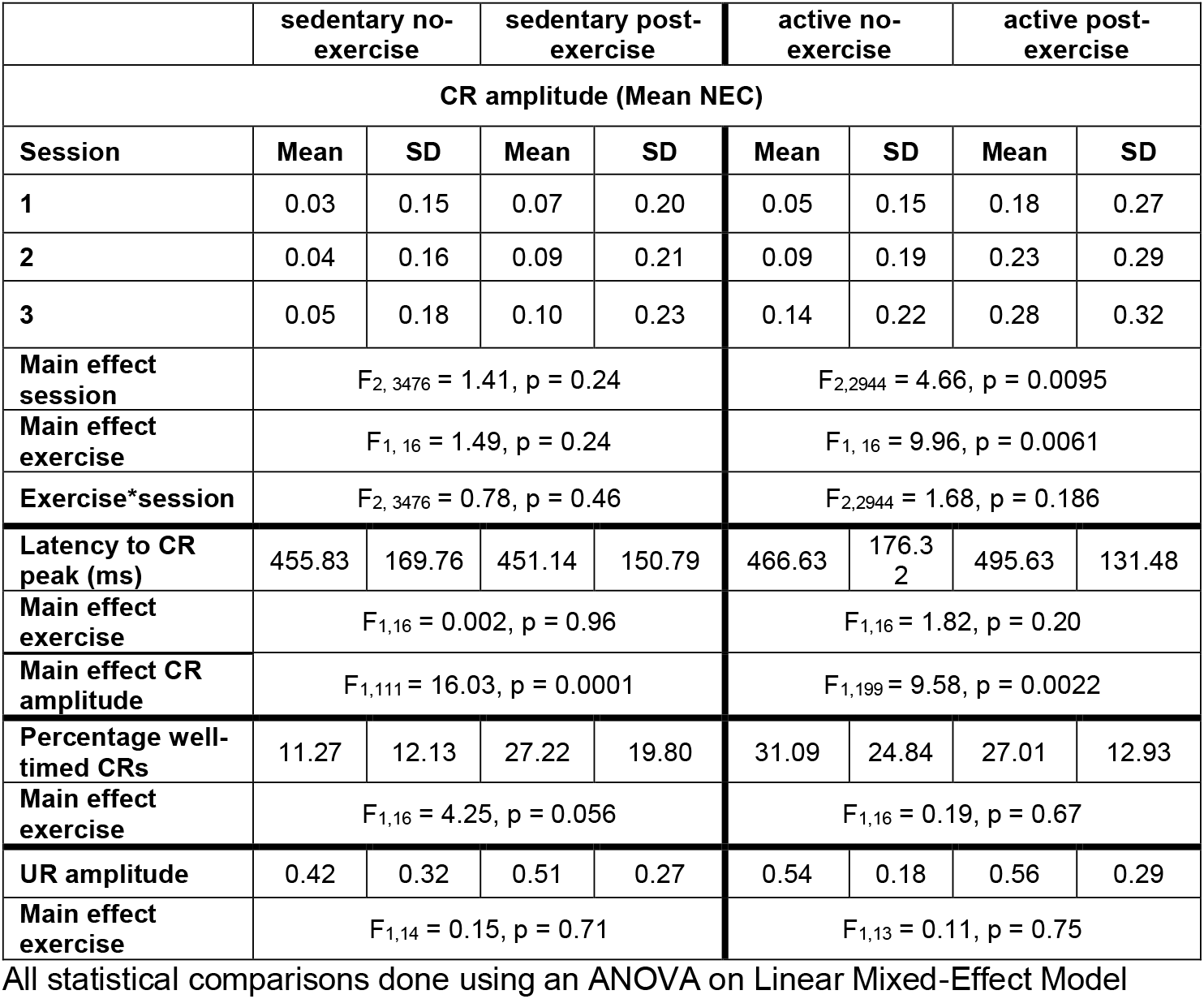
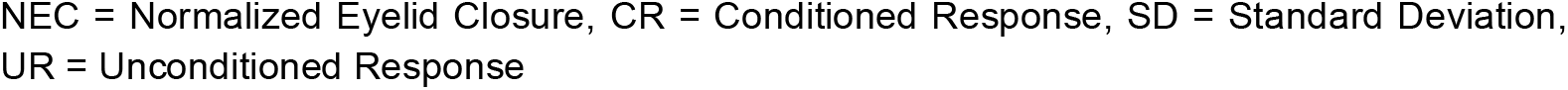
Conditioned response amplitudes and latencies in active and sedentary groups with and without an exercise intervention prior to eyeblink conditioning.

|  | sedentary no-exercise |  | sedentary post-exercise |  | active no-exercise |  | active post-exercise |  |
| --- | --- | --- | --- | --- | --- | --- | --- | --- |
| CR amplitude (Mean NEC) |  |  |  |  |  |  |  |  |
| Session | Mean | SD | Mean | SD | Mean | SD | Mean | SD |
| 1 | 0.03 | 0.15 | 0.07 | 0.20 | 0.05 | 0.15 | 0.18 | 0.27 |
| 2 | 0.04 | 0.16 | 0.09 | 0.21 | 0.09 | 0.19 | 0.23 | 0.29 |
| 3 | 0.05 | 0.18 | 0.10 | 0.23 | 0.14 | 0.22 | 0.28 | 0.32 |
| Main effect session | F<sub>2, 3476</sub> = 1.41, p = 0.24 |  |  |  | F<sub>2,2944</sub> = 4.66, p = 0.0095 |  |  |  |
| Main effect exercise | F<sub>1, 16</sub> = 1.49, p = 0.24 |  |  |  | F<sub>1, 16</sub> = 9.96, p = 0.0061 |  |  |  |
| Exercise*session | F<sub>2, 3476</sub> = 0.78, p = 0.46 |  |  |  | F<sub>2,2944</sub> = 1.68, p = 0.186 |  |  |  |
| Latency to CR peak (ms) | 455.83 | 169.76 | 451.14 | 150.79 | 466.63 | 176.32 | 495.63 | 131.48 |
| Main effect exercise | F<sub>1,16</sub> = 0.002, p = 0.96 |  |  |  | F<sub>1,16</sub> = 1.82, p = 0.20 |  |  |  |
| Main effect CR amplitude | F<sub>1,111</sub> = 16.03, p = 0.0001 |  |  |  | F<sub>1,199</sub> = 9.58, p = 0.0022 |  |  |  |
| Percentage well-timed CRs | 11.27 | 12.13 | 27.22 | 19.80 | 31.09 | 24.84 | 27.01 | 12.93 |
| Main effect exercise | F<sub>1,16</sub> = 4.25, p = 0.056 |  |  |  | F<sub>1,16</sub> = 0.19, p = 0.67 |  |  |  |
| UR amplitude | 0.42 | 0.32 | 0.51 | 0.27 | 0.54 | 0.18 | 0.56 | 0.29 |
| Main effect exercise | F<sub>1,14</sub> = 0.15, p = 0.71 |  |  |  | F<sub>1,13</sub> = 0.11, p = 0.75 |  |  |  |

All statistical comparisons done using an ANOVA on Linear Mixed-Effect Model
NEC = Normalized Eyelid Closure, CR = Conditioned Response, SD = Standard Deviation, UR = Unconditioned Response

In contrast, in the active group the CR amplitude differed between the no and post-exercise interventions (**figure 1D, F, table 2**). In the no exercise intervention, the mean CR amplitude at session 1 was 0.05 (±0.15) and increased to 0.14 (±0.22) by session 3. The mean CR amplitude for the post-exercise intervention was higher than the no exercise intervention at session 1 (mean=0.18 ±0.27), increasing to 0.28 (±0.32) by session 3. In the active group, the effect of ‘exercise’ (F_1,16_=9.96, p=0.0061) and ‘session’ (F_2,2944_=4.66, p=0.0095) were significant. The interaction between ‘exercise’ and ‘session’ was not significant (F_2,2944_=1.68, p=0.19). Post-hoc tests showed a significant difference between the active no and post-exercise interventions at sessions 1 (t_16_=-2.73, p=0.029) and 2 (t_16_=-3.36, p=0.012; **supplemental table 1**). In the active group with no exercise intervention, post-hoc tests showed a significant difference in CR amplitude between sessions 1 and 3 (t_2944_=-1.98, p = 0.048). Likewise, in the active group with the exercise intervention, post-hoc tests showed a significant difference in CR amplitude between sessions 1 and 3 (t_2944_=-2.39, p = 0.017)

### 3.4 Conditioning – timing

Next it was determined whether acute aerobic exercise had an effect on the latency to CR peak. For this, CS-only trials were analyzed. For both lifestyle groups, the CR peak times are roughly distributed around the expected onset of the US regardless of the intervention (**figure 2A, B**). Most of the variation in CR peak times could be explained by CR amplitude for both the sedentary (F_1,111_=16.03, p=0.0001) and active (F_1,199_=9.58, p=0.0022) lifestyle groups. The latencies to CR peaks did not differ significantly between no-exercise and post-exercise interventions for both sedentary (F_1,16_=0.002, p= 0.96) and active (F_1,16_=1.82, p=0.20, **table 2)** individuals.

**Figure 2.**
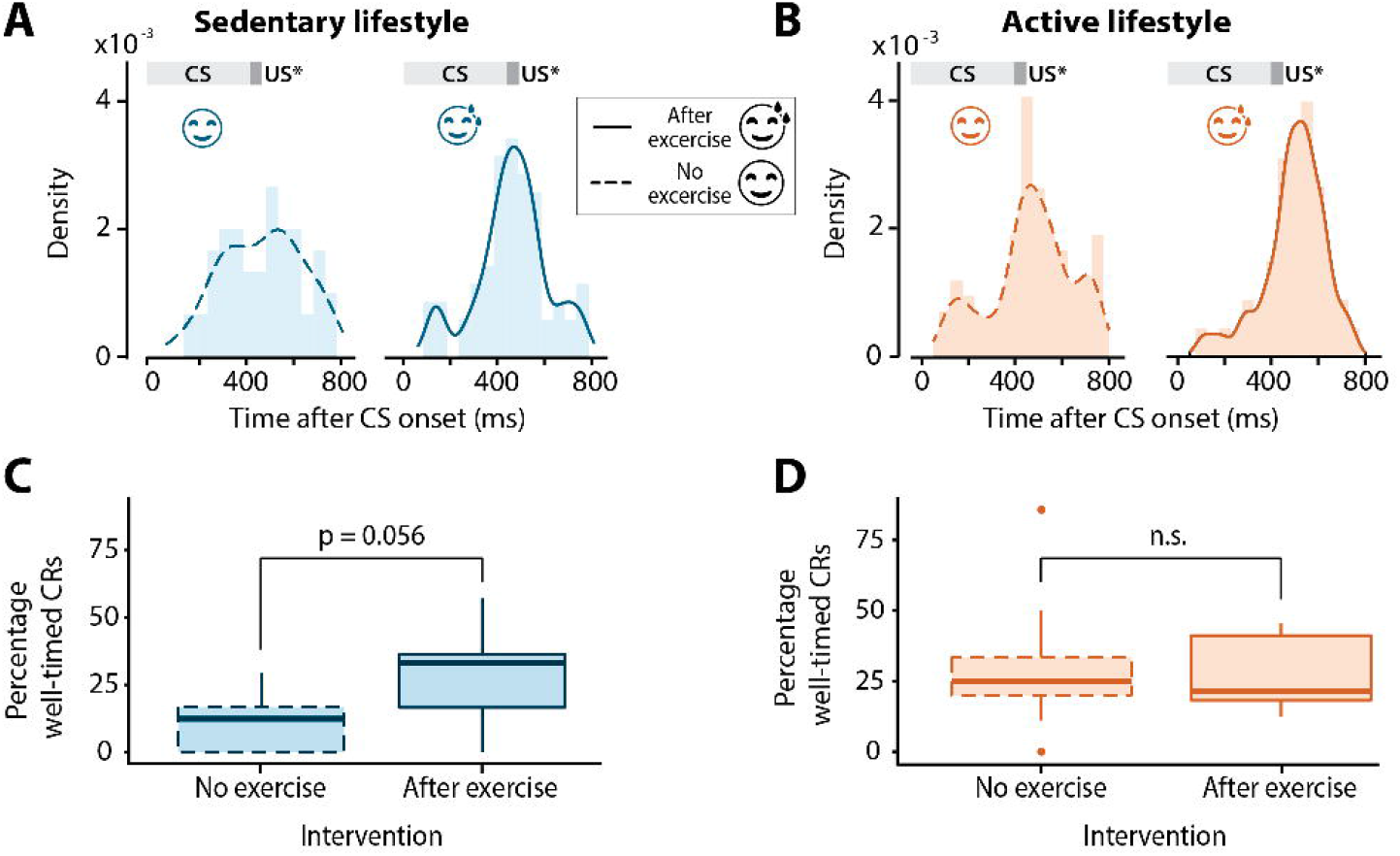
Timing of conditioned eyeblink responses in sedentary and active individuals with or without exercise. **(A)** Distribution of latency to conditioned response peak for all conditioned stimulus (CS) only trials across all sessions in sedentary and **(B)** active groups with or without exercise preceding eyeblink conditioning sessions. The dark grey block at 400 ms indicates the expected onset of the unconditioned stimulus (US*) which is omitted in these trials. The light grey block indicates the presentation of the CS. Note the distribution centered roughly around the expected onset of the US at 400 ms for all groups. **(C)** Boxplots of percentage of well-timed conditioned responses (CRs) in the sedentary and **(D)** active groups with or without exercise. Middle line indicates group medians, box ends indicate lower and upper quartiles, whiskers indicate group minima and maxima and dots indicate outliers. n.s. = not significant.

The mean percentage of well-timed CRs was also determined for active and sedentary lifestyle groups with and without the exercise intervention (**table 2, figure 2C,D)**. While the percentage of well-timed CRs was quite low for the sedentary group with no exercise intervention (**figure 2C**, mean=11.27% ±12.13), the effect of exercise on the percentage of well-timed CRs was neither significant for the sedentary (F_1,16_=4.25, p=0.056) nor the active group (F_1,16_=0.19, p=0.67).

### 3.5 Unconditioned responses

To determine if the effect of aerobic exercise was specific to CRs or more generalized, unconditioned response amplitude was compared between interventions within lifestyle groups. The block 1 trials of session 1 were used to determine unconditioned response amplitude, as these data were obtained before onset of the CRs that could in principle influence the amplitude of the unconditioned response(21). In both the sedentary (F_1,14_=0.15, p=0.71) and active groups (F_1,13_=0.11, p=0.75), there was no significant difference in unconditioned response amplitude between the no and post-exercise interventions (**figure 3A, B, table 2)**.

**Figure 3.**
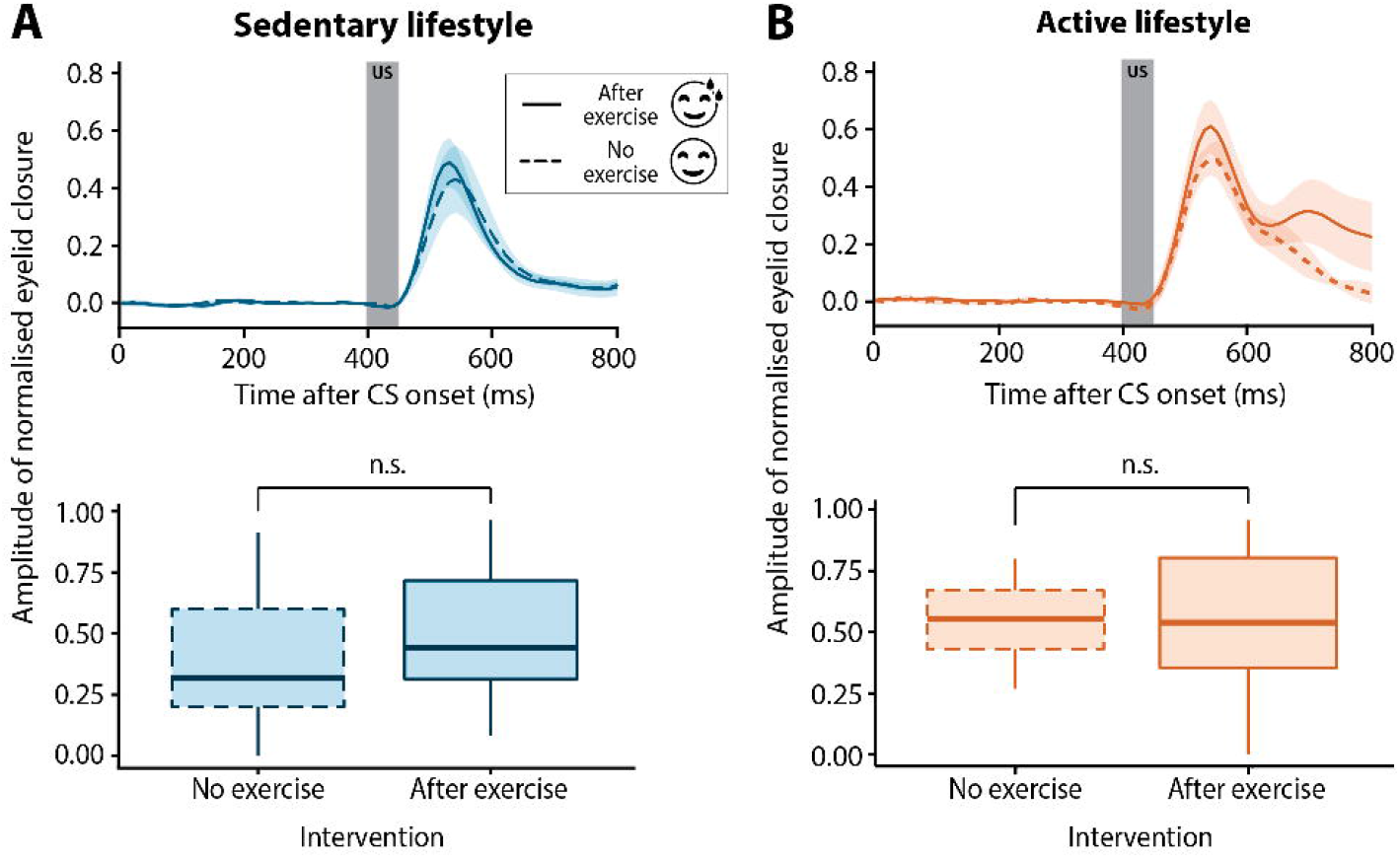
Unconditioned stimulus auditory-evoked blinks in sedentary and active groups with or without exercise. **(A)** Group averaged unconditioned response amplitudes for sedentary or **(B)** active individuals with (solid line) or without (dashed line) exercise preceding the eyeblink conditioning session. Unconditioned response amplitudes were calculated for the first two blocks of session 1, prior to the development of conditioned responses. The dark grey block (top panels) from 400 - 450 ms indicates the presentation of the unconditioned stimulus (US). The unconditioned response amplitude was similar regardless of the exercise intervention for both sedentary and active groups. In the boxplots (bottom panels), the middle line indicates group medians, box ends indicate lower and upper quartiles, whiskers indicate group minima and maxima. n.s. = not significant.

## 4 DISCUSSION

Aerobic exercise enhanced CR acquisition in a smartphone-mediated eyeblink conditioning paradigm. This effect of exercise was, however, only seen in individuals with an active lifestyle. This finding parallels that of Hopkins and colleagues(22), where acute exercise enhanced recognition memory in individuals with a prior four week exercise training program, but not in individuals without such a program. Exercise had no major effect on the unconditioned response amplitude or the timing of CR peaks.

### 4.1 Conditioned response acquisition

Both the active post-exercise and no-exercise interventions showed a significant increase in CR amplitude over the three sessions. The significant differences in CR amplitude between the active no-exercise and active post-exercise groups at early sessions parallels the acute exercise enhancing effect seen in animal research where mice running at faster speeds showed CRs in earlier sessions compared to mice running at slower speeds(8). Exercise may have a priming effect; reducing the number of practice sessions needed for implicit learning(23). This enhancing effect of exercise was specific to CRs as the amplitude of the unconditioned responses did not differ between groups. It is proposed that locomotor activity acts directly within the cerebellar cortex to modulate eyeblink conditioning. Locomotor activity signaling via cerebellar mossy fibers (MF) may converge with the CS MF signaling hereby facilitating learning(8). While exercise may have acted directly within the cerebellar cortex to enhance associative learning, it is unclear why such an effect would differ for active and sedentary individuals.

The finding that acute exercise facilitates eyeblink conditioning in active but not sedentary individuals may point towards a mechanistic role of neuropeptidergic transmitters and/or neurotrophins. Indeed, both human(22,24,25) and animal(26) studies on neuropeptidergic transmitters and neurotrophins show differential effects of acute exercise in active compared to sedentary subjects. Likewise, the dopaminergic, adrenergic and norepinephrinergic pathways, which are all catecholaminergic systems that prominently co-release neuropeptides, are upregulated in humans(2,27) and animals(28–30) following exercise. While the proposed role of these neurotransmitters in exercise-induced cognitive benefits are frequently studied(2,31), their potential influence on associative learning has received less attention(27,32). Despite this, there is evidence for a role of neurotransmitters in cerebellar learning. In rabbits, pharmacological monoamine depletion resulted in a dose-dependent reduction in CRs in an eyeblink conditioning task(33). Additionally, in rats, cerebellar norepinephrine was shown to be involved in the acquisition of CRs(34,35). These findings may extend to humans, where increased levels of norepinephrine following exercise have been associated with improved motor skill acquisition compared to resting controls(27) and where chronic training increased the plasma catecholamine response, compared to no training, after a cycling task(36).

Similarly, the neurotrophin BDNF may facilitate exercise-induced brain plasticity(26,37,38) and memory formation(3,39). Notably, BDNF mutant mice show impaired eyeblink conditioning(40) and the impact of tDCS on eyeblink conditioning in humans can depend on BDNF mutations(41). Moreover, a meta-analysis of the effects of exercise on BDNF in humans reported an enhanced BDNF response to acute exercise in active compared to sedentary individuals(25). Thus, it is possible that in this study the acute exercise in the sedentary group was not sufficient to induce the BDNF levels needed to enhance eyeblink conditioning. Together, these findings provide tentative molecular clues as to why in this study aerobic exercise enhanced associative learning in active but not sedentary individuals.

### 4.2 Conditioned response timing

Unlike the acquisition of CRs, the timing of these responses did not significantly differ across groups. Similarly, CR timing was unaltered in rats with access to a running wheel despite these rats showing enhanced CRs compared to non-exercising rats(21). Interestingly, while not significantly different, the percentage of well-timed CRs was slightly higher in the sedentary post-exercise compared to the sedentary no-exercise group. This is in-line with a recent study where physically enriched mice showed more well-timed CRs compared to mice in standard environments(10) and may require further research in a larger sample.

### 4.3 Limitations and future work

Despite all participants being instructed to exercise at moderate intensity, the average self-reported exercise intensity rating differed between active and sedentary individuals, with sedentary individuals perceiving the exercise to be harder than active individuals. While not directly comparable to a physical stressor, a psychosocial stressor impaired eyeblink conditioning in human subjects(42). The increased perceived intensity of exercise in the sedentary group may have masked the exercise enhancing effects seen in the active group. Interestingly, however, the average heart rate during the intervention was not significantly different between lifestyle groups, although not all participants recorded their heart rate. Future work could further explore potential associations between exercise intensity and eyeblink conditioning using more objective measures of exercise intensity.

A second limitation of this study was the lack of power to analyze sex differences between groups. A sex-dependency in eyeblink conditioning performance has been shown, with females showing more CRs compared to males(13). The generalizability of the findings are limited as the study was conducted on mainly white participants with middle to high socio-economic status. As this was not a randomized controlled trial (RCT), direct causal relationships between exercise and associative learning cannot be made. A future RCT could assess the level of exercise training required before acute effects of exercise on learning are seen.

Finally, future directions could include investigating a possible clinical application of these findings. Studies have shown impaired eyeblink conditioning in various patient groups, for example schizophrenia(43), ADHD(44) and spinocerebellar ataxia(45). Whether exercise could restore eyeblink conditioning deficits in these patient groups and hereby inform exercise-based therapeutic interventions remains to be determined.

## 5 Conclusions

Acute aerobic exercise enhanced the acquisition of associative learning in an eyeblink conditioning paradigm for individuals with a regularly active lifestyle. These results tentatively confirm in humans what has been shown in animals regarding the facilitatory effects of exercise on eyeblink conditioning. By focusing on a well-characterized learning paradigm, this study contributes to a more objective understanding of how exercise influences the brain, however, given the exploratory nature of this study, replicating the results in a larger cohort will be an important next step.

## DECLARATIONS

### Data availability

Data are available from the corresponding authors upon reasonable request.

### Ethics approval

This study involved human participants and was approved by the Institutional Review Board for Human Subjects of Princeton University (IRB #13943). The study was conducted in line with the principles stated in the Declaration of Helsinki.

### Consent to participate

Participants gave written informed consent prior to taking part in this study

### Consent for publication

Consent for publication of the image used in Figure 1B was obtained from the relevant individual.

### Author contributions

KDG and HJB conceived and designed this study. KDG, HJB, CPB, SKEK and LEMR were responsible for the methodology. KDG carried out the investigation. KDG, HJB and SKEK carried out the analysis with technical assistance provided by CPB. Visualisation was done by HJB, KDG and SKEK. HJB, CIDZ and SKEK were responsible for supervision. The original draft of the manuscript was written by KDG and reviewing and editing was done by HJB, CIDZ, SKEK, CPB, LEMR and ES. All authors read and approved the final version of the manuscript.

## Acknowledgements

We would like to thank all participants for their valuable cooperation in this study.

