## supplemental figure 1 for "Acute Aerobic Exercise Enhances Associative Learning in Active but not Sedentary Individuals"

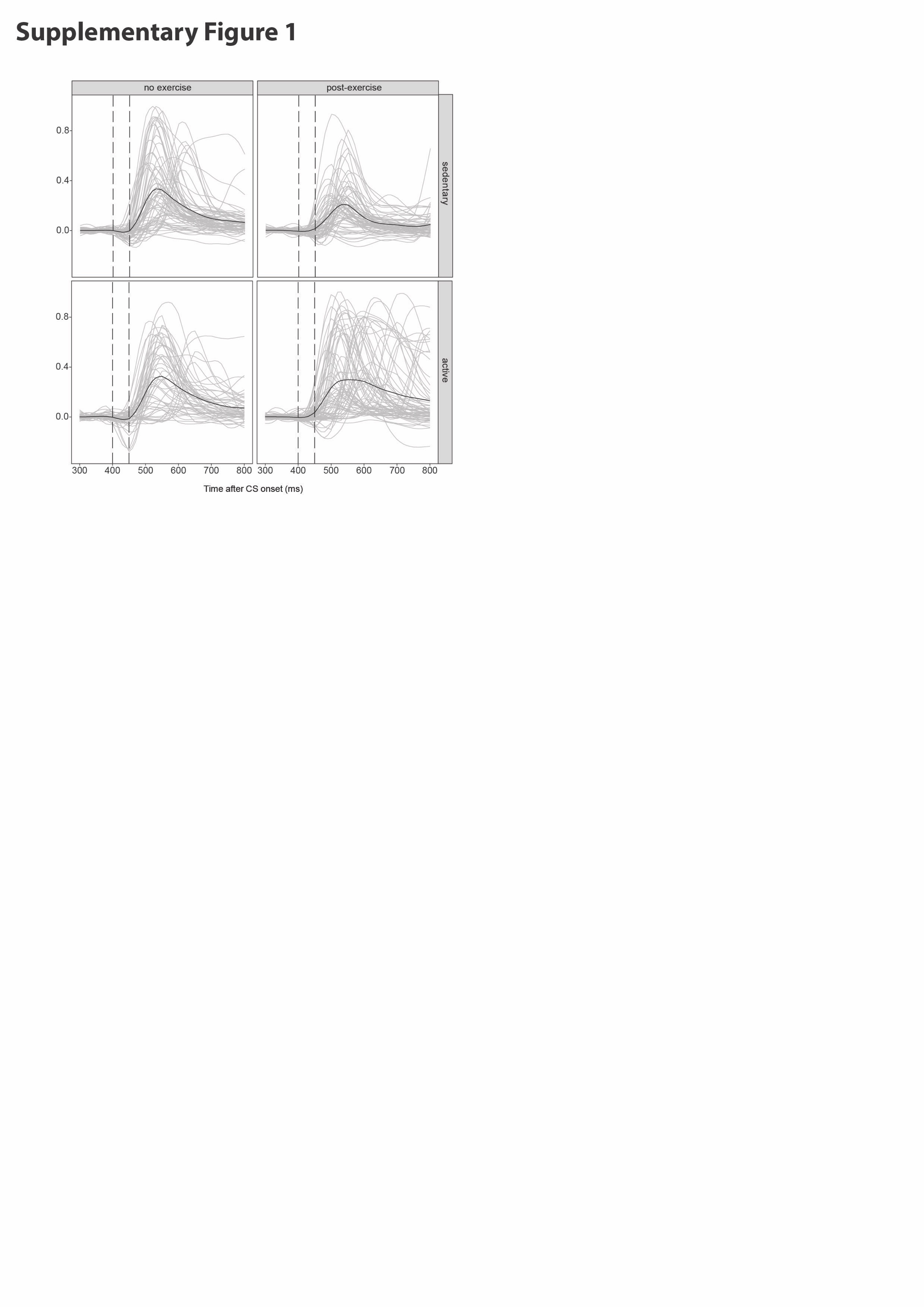

**Supplementary Figure 1 | Raw traces of auditory evoked blinks for sedentary and active groups with or without an exercise intervention.** Unconditioned stimulus only trials from session 1 in sedentary and active individuals without (left panels) and directly after (right panels) an exercise intervention. Grey traces represent subject averages and black traces represent group averages. The first vertical dashed line represents the onset of the unconditioned stimulus at 400 ms and the second dashed line represents the offset of the unconditioned stimulus at 450 ms. Note the latency in response to presenting the unconditioned stimulus in all groups.
