## supplemental figure 2 for "Acute Aerobic Exercise Enhances Associative Learning in Active but not Sedentary Individuals"

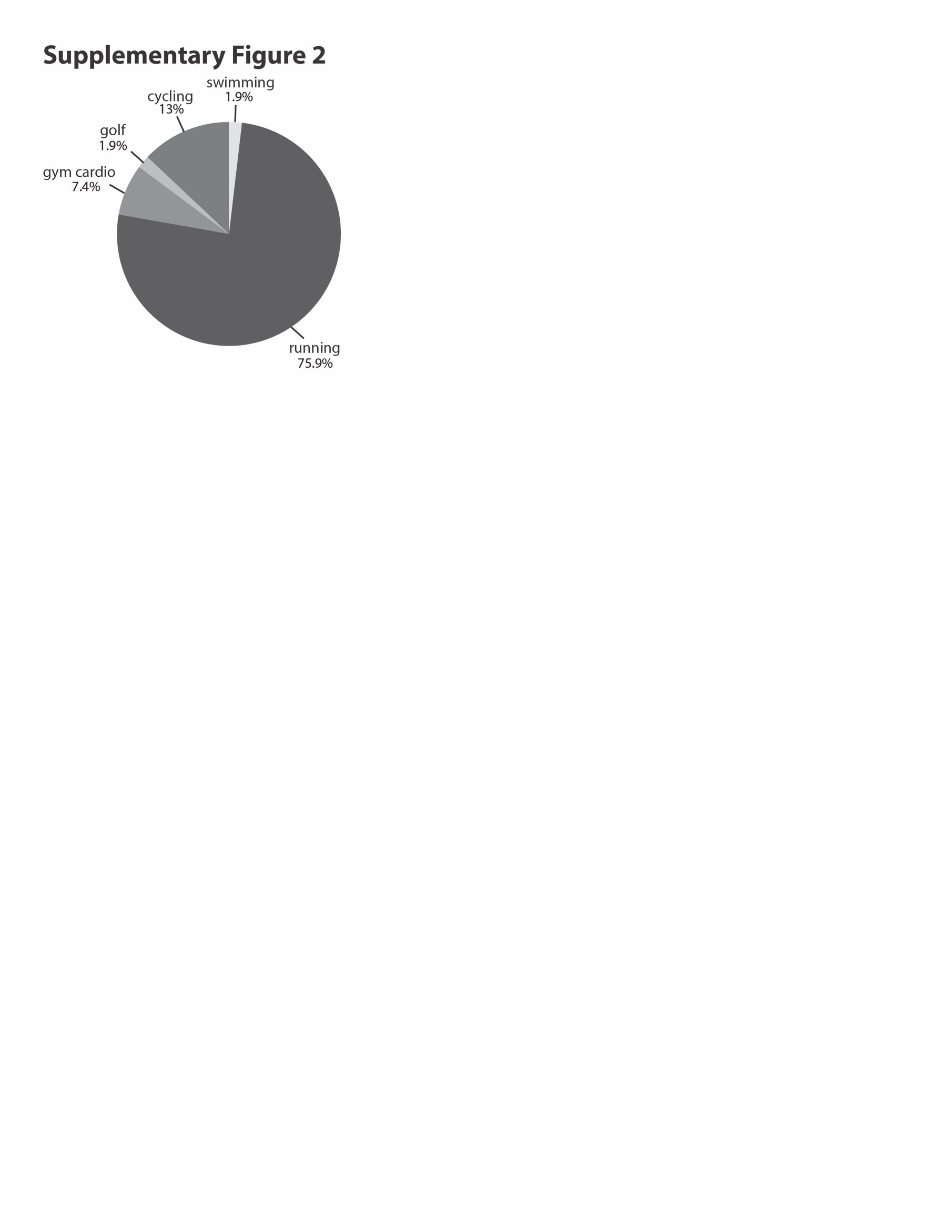

**Supplementary Figure 2** | **Frequency of types of exercise done prior to eyeblink conditioning in the exercise intervention**

Distribution of exercise activities done by active and sedentary individuals in the exercise intervention. Distribution includes all three exercise sessions per participant done prior to the three eyeblink conditioning sessions
