## supplemental figure 3 for "Acute Aerobic Exercise Enhances Associative Learning in Active but not Sedentary Individuals"

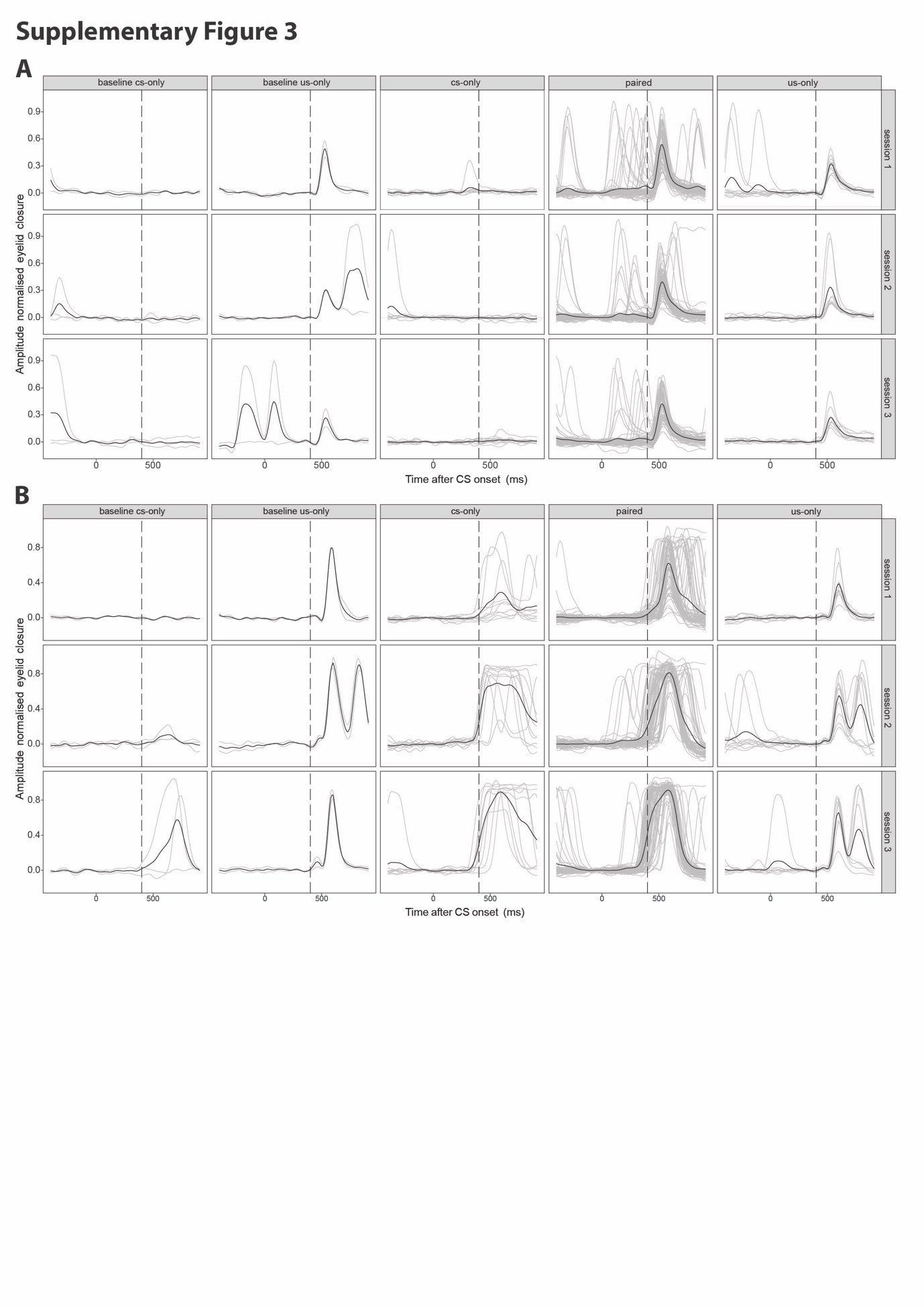

**Supplementary Figure 3** | **Raw eyeblink traces from two participants for all trial types.** Example traces of a poor learner, from the sedentary no-exercise intervention **(A)** and a learner from the active post-exercise intervention **(B)** for all trials over 3 sessions of delay eyeblink conditioning. Vertical line at 400 ms showing expected (conditioned stimulus (CS) -only trials) or actual presentation of the unconditioned stimulus (US). Gray traces indicate trials, black traces indicate session averages. Numbers on the right hand side indicate session numbers.
