## supplemental table 2 for "Acute Aerobic Exercise Enhances Associative Learning in Active but not Sedentary Individuals"

**Supplemental Table 1 | Post-hoc tests within sedentary and active groups for no vs. post-exercise interventions prior to three sessions of eyeblink conditioning**

|  | **Pairwise differences** | | | |
| --- | --- | --- | --- | --- |
|  | **estimate** | **t-ratio** | **p-value** | **adjusted p-value** |
| **Sedentary** |  | | | |
| Session 1: no vs. post-exercise | -0.19 | -0.98 | 0.34 | 0.68 |
| Session 2: no vs. post-exercise | -0.28 | -1.43 | 0.17 | 0.52 |
| Session 3: no vs. post-exercise | -0.22 | -0.98 | 0.34 | 0.68 |
| Sedentary no exercise: session 1 vs. session 3 | -0.14 | -1.12 | 0.26 | 0.26 |
| Sedentary post-exercise: session 1 vs. session 3 | -0.16 | -1.30 | 0.20 | 0.20 |
| **Active** |  | | | |
| Session 1: no vs. post-exercise | -0.44 | -2.73 | 0.014 | 0.029 |
| Session 2: no vs. post-exercise | -0.60 | -3.36 | 0.0040 | 0.012 |
| Session 3: no vs. post-exercise | -0.48 | -1.99 | 0.064 | 0.064 |
| Active no exercise: session 1 vs. session 3 | -0.31 | -1.98 | 0.048 | 0.048 |
| Active post-exercise: session 1 vs. session 3 | -0.35 | -2.39 | 0.017 | 0.017 |

All statistical comparisons done using an ANOVA on Linear Mixed-Effect Model

P-value adjustment: Bonferroni-Holm method for three tests for comparisons between interventions
